# *De novo* Transcriptomic Analysis to Understanding Piperine Biosynthesis in *Piper chaba* Hunter

**DOI:** 10.1101/2025.02.28.640931

**Authors:** Deeksha Singh, Rajiv Ranjan

## Abstract

Medicinal plants serve as invaluable sources of bioactive compounds, yet the molecular basis of their secondary metabolite biosynthesis remains largely unexplored. *Piper chaba* Hunter, an important but understudied member of the Piperaceae family, is known for its pharmacologically active alkaloids, particularly piperine. This study presents the first comprehensive transcriptomic analysis of *P. chaba* to uncover the genetic pathways regulating its metabolite production. The quantification of piperine, a major bioactive compound, was conducted using UPLC, which revealed its highest concentration in the spike (331.3 mg/g) followed by the stem (63.5 mg/g), root (10.3 mg/g), and leaves (2.82 mg/g). High-quality RNA sequencing of leaves, roots, and spikes using next-generation sequencing (NGS) generated 228,481 transcripts, and 184,574 unigenes were identified after redundancy removal. Coding sequences (CDS) derived from these unigenes were annotated using BLASTX and KEGG databases, which highlighted significant metabolic pathways, including those related to piperine biosynthesis. A total of thirteen genes associated with piperine biosynthesis were identified. Validation of nine selected genes, including Farnesyl pyrophosphate synthase and Piperic acid synthase, was performed through qRT-PCR using the 2^-ΔΔCt^ method, confirming their involvement in piperine and other metabolite biosynthesis. Functional annotation categorized the CDS into Gene Ontology domains, with transcription factors such as bHLH and NAC families playing prominent roles in metabolic regulation. Additionally, 5,050 SSRs were identified, offering potential markers for genetic studies. This pioneering study establishes a molecular framework for understanding the biosynthetic pathways of *P. chaba*, providing valuable insights for its application in sustainable medicine and agriculture.

## Introduction

Herbal medicines have been essential to human health and survival since ancient times, serving as the foundation of early healthcare systems (Qadir and Raja, 2021). The transition from conventional medicine to modern pharmacology has primarily been accelerated by the investigation of medicinal plants and their phytoconstituents. Among these, the genus *Piper* is of particular significance within the family Piperaceae, comprising approximately 1,000 species found in tropical and subtropical regions (Durant-Archibold et al. 2018). Several species within this genus, including *Piper nigrum* (black pepper), *Piper betle* (betel leaf), and *Piper longum* (long pepper), have been extensively studied for their bioactive constituents and therapeutic applications.

Beyond these well-known species, the *Piper chaba* Hunter represents a lesser-explored yet economically and pharmacologically significant species (Alam and Naser, 2020). *P. chaba* is a perennial flowering vine in the Piperaceae family. It is indigenous to South and Southeast Asian countries and commonly known as Badi Pippali or Bangla Tipali in India (Rameshkumar et al. 2011; Ruangnoo et al. 2012; Islam et al. 2020). Morphologically, *P. chaba* is a creeping vine that grows on the ground and can also climb trees. The plant contains oval leaves, often about 3 to 5 inches in length. The flowers are monoecious and zygomorphic. These flowers predominantly blossom in the monsoon season. The fruits are elongated, reaching up to 3 inches in length, turning red when ripe, and dark brown or black when dried (Ruangnoo et al. 2012; Islam et al. 2020).

Traditional medicinal applications of *P. chaba* primarily utilize its stem, root, and fruit, which contain several bioactive compounds, including alkaloids such as piperine, piperanine, pipernonaline, piperamine 2, 4-decadienoic acid piperidide, kusunokinin, and pellitorine ((Islam et al. 2020; Sarfaraz et al. 2014). The root of *P. chaba* possesses alexiteric properties, making it beneficial for treating asthma, bronchitis, and consumption. The fruit possesses stimulating and carminative qualities, making it beneficial for treating hemorrhoidal symptoms (Islam et al. 2021). The stem of this plant is utilized for alleviating post-delivery discomfort in mothers, and for reducing rheumatic pains and diarrhea (Taufiq-Ur-Rahman et al. 2005). The edible portion of the plant is the swollen stem, which is highly efficacious in combating colds and coughs, while also improving immunity against these diseases (Sarkar, 2012). Pharmacological investigations have revealed that *P. chaba* possesses antibacterial, carminative, expectorant, analgesic, hypotensive, and smooth muscle relaxant properties. Piperine, a potent alkaloid primarily found in *P. chaba* and other members of the *Piper* genus has a diverse array of pharmacological effects. These encompass antibacterial, immunomodulatory, anti-mutagenic, hepatoprotective, antioxidant, antimetastatic, and anticancer properties. Chabamide, a dimeric amide alkaloid extracted from the stem of *P. chaba,* has antimalarial, antituberculosis, and cytotoxic effects (Rukachaisirikul et al. 2002; Rao et al. 2011; Ren et al. 2015). Similarly, piplartine, an alkaloid amide derived from this plant, is another notable anticancer candidate (Martha Perez Gutierrez et al. 2013; Vadaparthi et al. 2018).

Despite its extensive ethnobotanical and pharmacological significance, the molecular mechanisms underlying the biosynthesis of its bioactive compounds remain unexplored. Transcriptomic studies provide crucial insights into gene expression patterns and regulatory pathways involved in secondary metabolite production, offering a foundation for further biotechnological and pharmaceutical applications (Tyagi et al. 2022; Tyagi et al. 2024; Singh et al. 2024). However, no such studies have been conducted on *P. chaba*, leaving a critical gap in scientific knowledge.

To address this, the present study aims to perform the first-ever transcriptomic analysis of *P. chaba*, elucidating key genes and pathways associated with its bioactive compounds. This research will enhance the scientific understanding of *P. chaba* and support its sustainable utilization in medicine, agriculture, and biopharmaceutical industries by uncovering the molecular basis of its pharmacologically active constituents. The findings will also provide a valuable genomic resource for future studies on metabolic engineering, conservation, and bioprospecting of this important medicinal plant.

## Material and methods

In January 2023, fresh and healthy plant samples of *Piper chaba,* including root, stem, leaves, and spike were collected from the Herbal Garden of Dayalbagh Educational Institute (DEI), Dayalbagh, Agra. The freshly collected healthy plant samples were carefully rinsed first with groundwater and subsequently, utilized distilled water to remove any adhering dirt. The samples were then air-dried in a shaded area at 25°C for 12-15 days to preserve the active compounds. Afterward, the dried plant materials were pulverized with a machine grinder. The pulverized samples were subjected to sieving using a 2 mm mesh sieve.

### (A) Quantitative analysis of the piperine by UPLC-PDA

- **Sample preparation** The 5 g of powdered *P. chaba* root, stem, leaves, and spike were refluxed individually in 50 ml of methanol for 6 hours at a steady temperature. Samples were concentrated using a rotary vacuum evaporator. A final concentration of 10 mg/ml was obtained by dissolving 10 mg of each sample in 1 ml of methanol. Following reconstitution, an aliquot of the solution was filtered using a 0.22 µm filter.
- **Standard solution preparation** A precisely measured amount (10 mg) of piperine was added to methanol (10 ml) to make a stock solution of 1 mg/ml. The solution was degassed in a sonicator for 10 minutes and subsequently filtered using a 0.22 µm nylon membrane before dilution preparation (working solution). A standard solution of piperine (1 mg/ml) was diluted into five distinct concentrations (25, 50, 100, 150 and 200 µg/ml). These solutions were subjected to the UPLC System for the preparation of a calibration graph. A calibration curve was formed by correlating the peak area with the piperine concentrations. The correlation coefficient consistently exceeded 0.997113 in all analyses (Chandra et al. 2015).
- **Chromatographic conditions and data interpretation** Analysis was done using an Acquity UPLC H-Class System (Waters, Milford, USA) with the quaternary solvent manager and PDA detector and a Hypersil Gold C18 column (2.1 mm x 100 mm, 1.9 mm particle size) (Acquity UPLC, Waters, Milford, USA) maintained at 25°C. Formic acid (0.1%) in water (buffer A) and formic acid (0.1%) in methanol (buffer B) were utilized as mobile phases. A flow rate of 0.200 ml/min was used, and a sample volume of 3μl was injected. Observations at 343 nm were made simultaneously. Piperine was quantified using a calibration curve (R² ≥ 0.997113) in the 25–200μg/ml range for all standards. The peak area of piperine in the sample extract was integrated and analyzed by comparing the calibration curve to evaluate the concentration. Software Empower 3 (Waters, Milford, USA) was utilized for processing chromatography data.

### (B) Transcriptomic analysis of different parts of the *P. chaba*

#### Plant material

In January 2023, fresh and healthy plant samples of *P. chaba* were collected from the Herbal Garden of Dayalbagh Educational Institute (DEI), Dayalbagh, Agra. Young leaves, roots, and spikes were carefully excised from healthy plants using a sterilized blade and immediately frozen in liquid nitrogen to preserve their integrity for further applications.

#### RNA isolation and library construction

Total RNA was obtained from three distinct tissues: leaves, root, and spike, utilizing the Alexgen Total RNA Extraction Kit. The paired-end sequencing libraries were constructed utilizing the KAPA mRNA HyperPrep Kit for Illumina as per the specified methodology. RNA was quantified using a Qubit® 4.0 fluorometer, while its quality was assessed by 1.2% agarose gel electrophoresis. mRNA enrichment was performed as per the user manual, followed by fragmentation of mRNA, synthesis of first and second-strand cDNA, end-repair, 3’ adenylation, adapter ligation, selective enrichment of adapter-ligated DNA fragments through PCR amplification, and subsequent validation of the library on the Agilent 4150 TapeStation. Cluster generation and sequencing were conducted using the Illumina Novaseq 6000 platform to produce 2×150 bp paired-end reads.

#### Transcriptome sequencing and analysis

- ***De novo* assembly (Master assembly)** High-quality adapter-trimmed reads from all *P. chaba* samples were combined to form a Master/Combine assembly using Trinity, with default parameters (k-mer size of 25). This produced a common assembly for the subsequent annotation of transcripts and investigation of differential expression across samples. The completeness of the *P. chaba* transcriptome was assessed using BUSCO analysis against the Viridiplantae_odb10 database (updated on December 5, 2024). Default parameters were applied to classify orthologs as complete, duplicated, fragmented, or missing, ensuring a standardized evaluation of transcriptome quality for downstream functional and comparative analyses.
- **Unigenes prediction from transcripts** Transcripts of combined assembly undergo additional processing for unigene prediction using the CD-HIT package. The CD-HIT-EST program was employed to eliminate shorter redundant transcripts that were completely covered by other transcripts showing >90% identity. The resultant non-redundant clustered transcripts were then categorized as unigenes.
- **CDS prediction from unigenes** CDS was anticipated from the unigene sequences utilizing TransDecoder software with default settings, indicating a minimum encoded protein length of 60 amino acids.
- **Transcription factor analysis** All CDS were searched against the transcription factor database (http://planttfdb.cbi.pku.edu.cn/download.php) using BLASTX with an e-value threshold of 1e-5 for the identification of transcription factor families.
- **Functional annotation and gene ontology (GO) analysis of predicted CDS** Functional annotation of the predicted coding sequences (CDS) of *P. chaba* was conducted by aligning them to non-redundant (NR) protein databases from the National Centre for Biotechnology Information (NCBI) using the BLASTX, with a minimum e-value threshold of 1e-5. BLAST 2.3.13+ software was employed for independent functional annotation and classification. All CDS exhibited BLASTX hits against the NR database on NCBI. Predicted CDS was also searched against KOG, Pfam, and Uniport databases using BLASTX. The optimal match output from each BLAST was utilized for annotating the CDS. After each BLAST search, annotation tags, irrespective of hits and including those with predicted annotations, were extracted from the following sequential BLAST search. The expected functions of CDS/proteins were elucidated through the use of gene ontology (GO) assignments. Gene ontology mappings provide a set of descriptive terms that characterize the qualities of gene products. The terms were categorized into three main domains: Biological Process (BP), Molecular Function (MF), and Cellular Component (CC). Gene Ontology (GO) mapping was conducted to obtain GO terms for all functionally annotated coding sequences (CDS) from BLASTX against the non-redundant database using OmixBox.
- **Classification of assembled transcriptomes** The KAAS server (http://www.genome.jp/kegg/ko.html) was utilized to functionally annotate the coding sequences (CDS) through BLAST comparisons with the KEGG gene database. The BBH (Bi-directional best hit) method was utilized to assign KEGG Orthology (KO) terms. The KEGG Orthology database was utilized for pathway mapping. The CDS were assignments of polypeptides produced by the combined assembly and mapped onto metabolic pathways according to KEGG.
- **Identification of Simple Sequence Repeats (SSRs)** SSRs, also known as microsatellites, are tandem repeats of nucleotide motifs ranging from 2 to 6 base pairs in length. They exhibit significant polymorphism and are universally found in all known genomes. Consequently, SSRs were discovered in the CDS sequences of each sample using the MISA Perl script. Identification was conducted using the following criteria: di-nucleotide patterns must occur a minimum of six times, while tri-, tetra-, penta-, and hexa-nucleotide patterns must appear at least five times.

#### Quantitative gene expression analysis of *P. chaba*

This study focused on the quantitative gene expression analysis of *P. chaba*, particularly genes involved in the piperine biosynthesis pathway. RNA was isolated using the PureLink® RNA mini kit and processed following standard protocols to ensure purity and integrity, which was confirmed via gel electrophoresis. First-strand cDNA was synthesized using the R2D 1st strand cDNA synthesis kit, with random or gene-specific primers. Primers for target genes, selected based on differential expression analysis, were designed using the PrimerQuest tool and optimized for qRT-PCR analysis. A total of nine genes, including an internal control (Ubiquitin C), were analyzed across leaves, roots, and spikes of *P. chaba* using the Brilliant III Ultra-Fast SYBR® Green QPCR kit. The qRT-PCR reactions were conducted in 96-well plates. Relative expression levels were calculated using the 2^-ΔΔCt^ method. Data from three biological replicates were statistically analyzed to identify significant expression patterns, enhancing the understanding of secondary metabolite biosynthesis in *P. chaba*.

## Results

### UPLC-based estimation of piperine content in different parts of the plant

The piperine content of *P. chaba* samples including root, stem, leaves, and spike was quantified using a UPLC method. Piperine concentrations were determined using the calibration curve. The calibration curve demonstrated that the computed piperine concentrations closely align with the anticipated values, signifying a strong correlation with negligible variance. All samples exhibited a predominant peak corresponding to piperine, as determined by an authentic piperine standard. A standard curve was established utilizing different concentrations of piperine (25–200 µg/ml), resulting in a correlation coefficient of 0.998556.

The quantification of piperine in various parts of *P. chaba* is presented in **Table 1**. The UPLC chromatograms of all samples, depicted in **Figure 1**, exhibited a consistent piperine profile throughout. Piperine was identified as the predominant compound in the spike, exhibiting a retention time of 0.800 minutes, comprising 100.00% of the area and a concentration of 331.3 mg/g. The stem demonstrated a retention time of 0.789 minutes, with piperine constituting 87.07% of the area and yielding 63.5 mg/g. The retention time of piperine in the root is 0.794 minutes, with 10.3 mg/g of piperine accounting for 75.06% of the area. The leaves exhibited minimal piperine concentration, with a retention time of 0.791 minutes, encompassing 19.17% of the area and with a total 2.82 mg/g piperine content.

**Figure 1:**
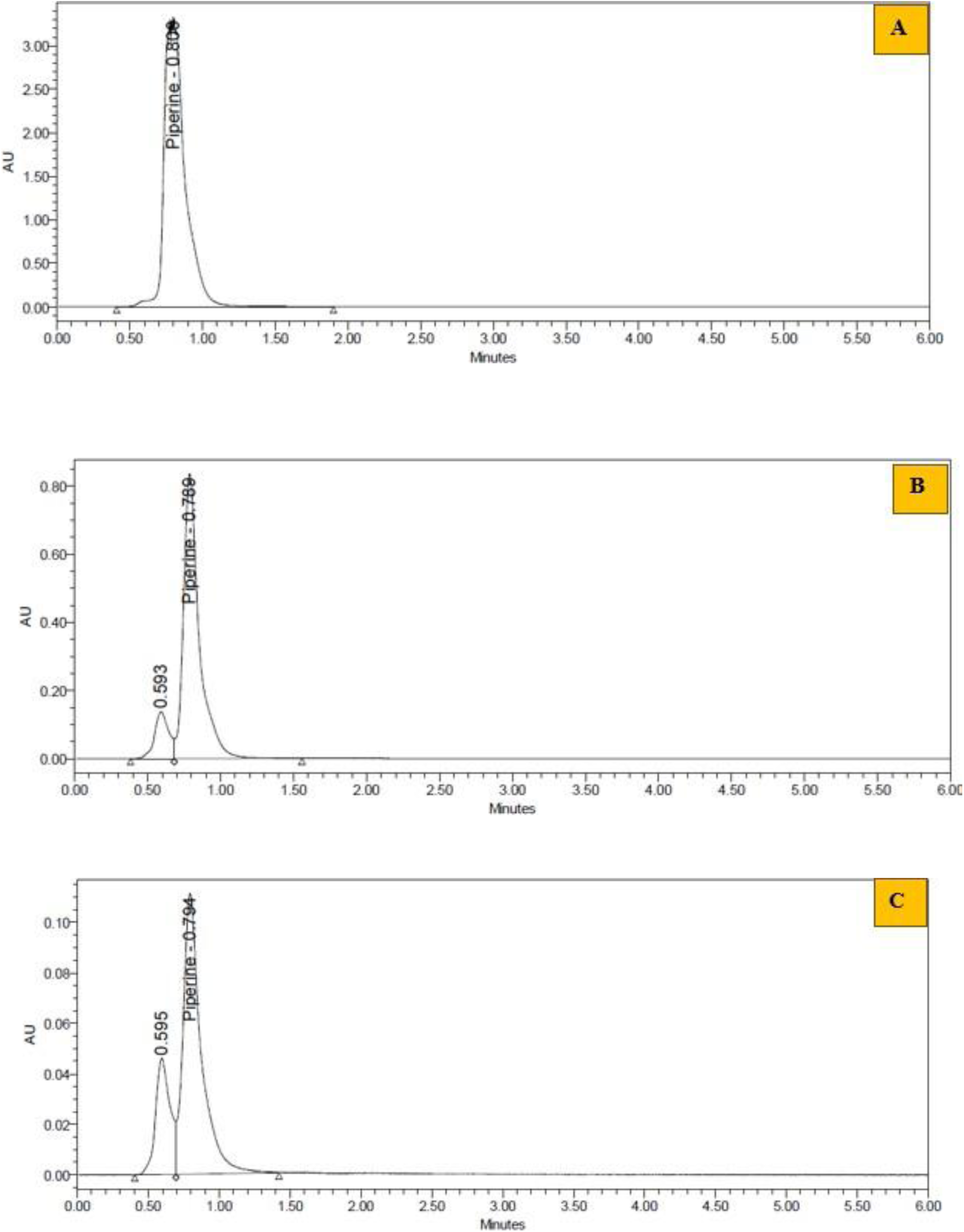
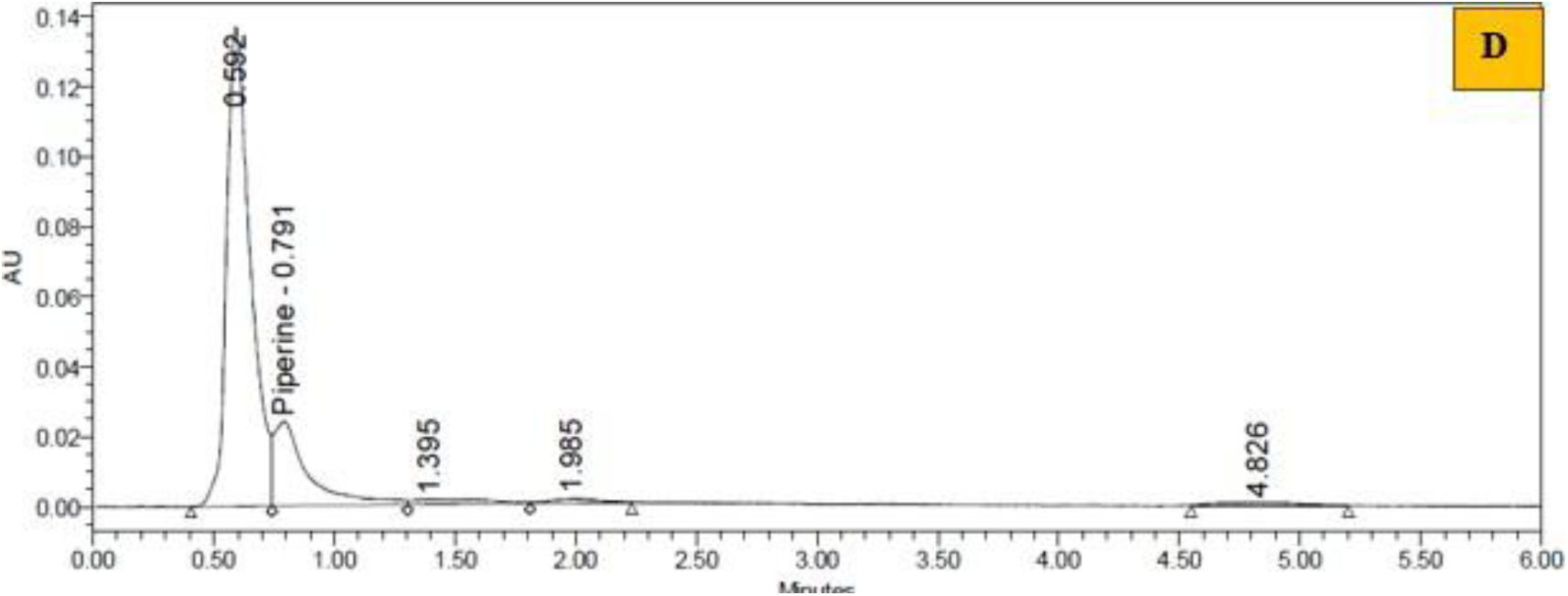
UPLC chromatograms of methanol extracts from different parts of *Piper chaba* (A) Spike, (B) Stem, (C) Root, (D) Leaves.

**Table 1:** UPLC-based quantification of piperine in different plant parts of *Piper chaba*.

| S. No. | Plant Part | Analyte | Retention Time (min) | Formula | % Area | Amount (mg/g) |
| --- | --- | --- | --- | --- | --- | --- |
| 1. | Spike | Piperine | 0.800 | C<sub>17</sub>H<sub>19</sub>NO<sub>3</sub> | 100.00 | 331.3 |
| 2. | Stem | Piperine | 0.789 | C<sub>17</sub>H<sub>19</sub>NO<sub>3</sub> | 87.07 | 63.5 |
| 3. | Root | Piperine | 0.794 | C<sub>17</sub>H<sub>19</sub>NO<sub>3</sub> | 75.06 | 10.3 |
| 4. | Leaves | Piperine | 0.791 | C<sub>17</sub>H<sub>19</sub>NO<sub>3</sub> | 19.17 | 2.82 |

### Transcriptome sequencing and *De novo a*ssembly

Next-generation sequencing (NGS) of *P. chaba* leaves, spike, and root was conducted using the Illumina Novaseq 6000 platform, yielding high-quality data through 2×150 bp paired-end (PE) reads. The high-quality reads from *P. chaba* leaves included 423,985,512 paired reads, the spike contained 663,405,588 paired reads, and the root had 703,265,582 paired reads. The *de novo* master assembly of high-quality adapter-trimmed paired-end reads from the sample generated 228,481 transcripts, as indicated in **Table 2**. The total transcriptome size was 25,068,437 bases. The average, maximum, and minimum transcript lengths in the libraries were 1097, 19061, and 300 bases, respectively. The length distribution of the raw reads and assembled transcripts from the libraries were characterized, and the data is presented in **Figure 2**. The BUSCO analysis of the *P. chaba* transcriptome demonstrated a high level of completeness, with 94.3% of BUSCO genes identified as complete (C). Among these, 8.2% were single-copy (S), while 86.1% were duplicated (D), indicating a considerable level of gene duplication. Additionally, 5.4% of the BUSCO genes were classified as fragmented (F), and only 0.3% were missing (M) (**Table 3**). The analysis was performed against a total of 425 BUSCO groups, confirming the high assembly quality and completeness of the *P. chaba* transcriptome. These findings suggest that the transcriptomic dataset provides a reliable foundation for downstream functional and comparative genomic analyses.

**Figure 2:**
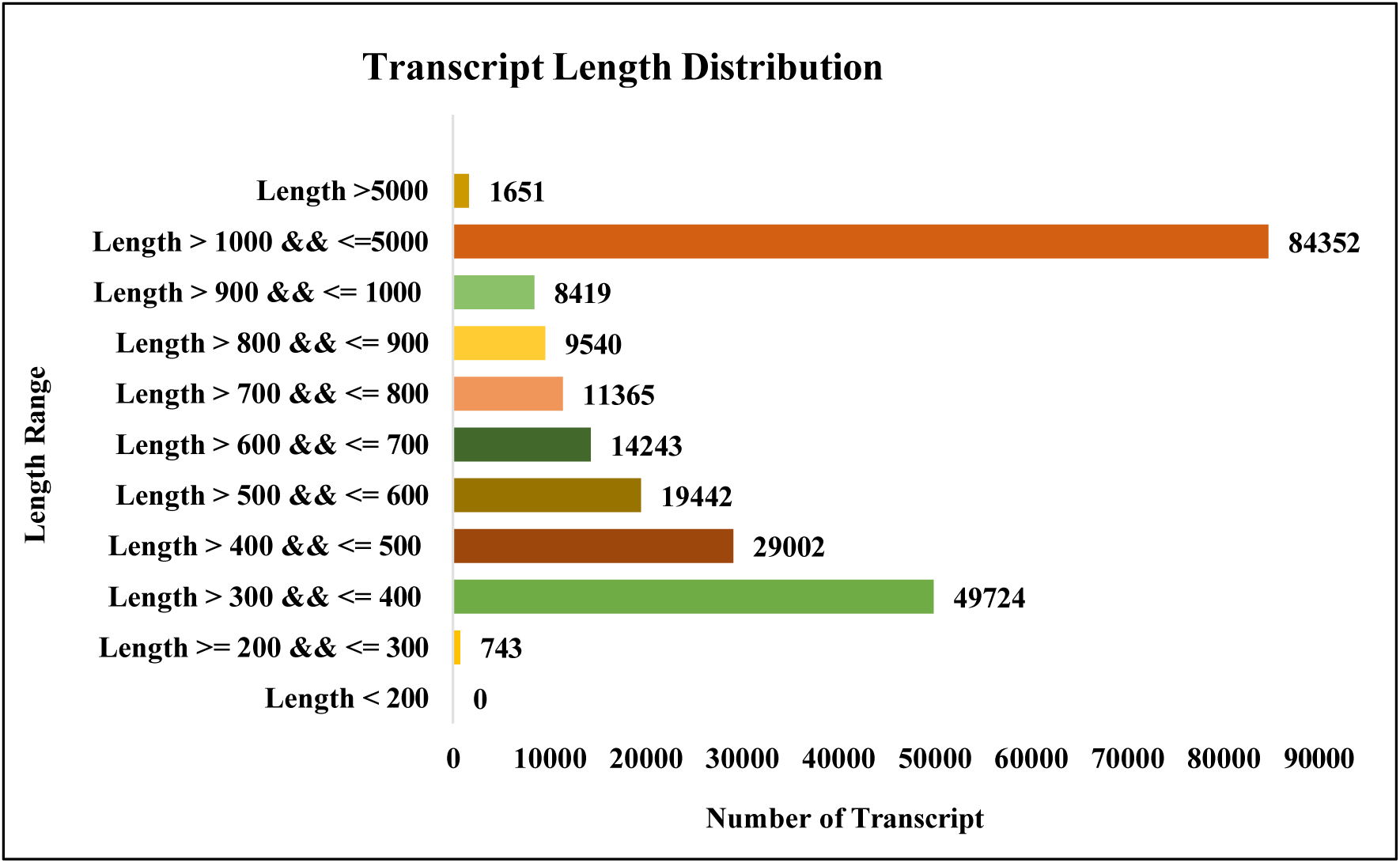
Length distribution of assembled transcripts. The highest number of coding sequences (CDS) were observed in the length range of over 1000 to 5000 base pairs, followed by the length range of over 300 to 400 base pairs.

**Table 2:**
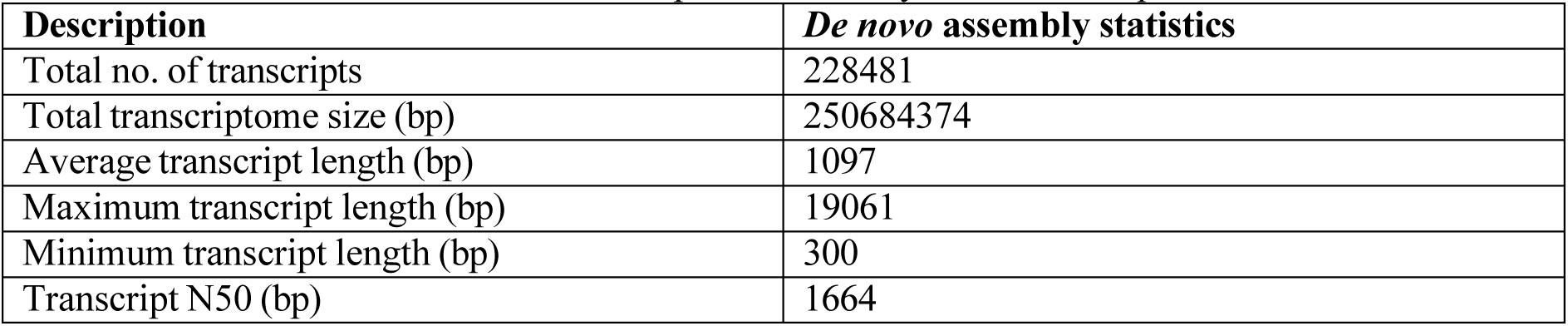
*De Novo* transcriptome assembly statistics of *Piper chaba*.

**Table 3:**
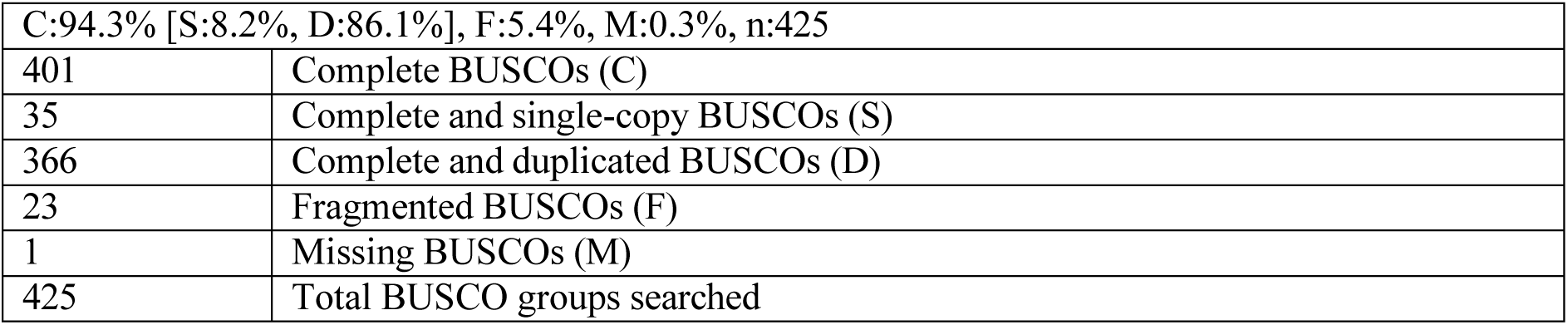
BUSCO analysis for *Piper chaba* transcriptome.

|  |  |
| --- | --- |
| C:94.3% [S:8.2%, D:86.1%], F:5.4%, M:0.3%, n:425 |  |
| 401 | Complete BUSCOs (C) |
| 35 | Complete and single-copy BUSCOs (S) |
| 366 | Complete and duplicated BUSCOs (D) |
| 23 | Fragmented BUSCOs (F) |
| 1 | Missing BUSCOs (M) |
| 425 | Total BUSCO groups searched |

### Unigenes prediction from master assembly

The transcripts underwent additional processing for unigene prediction utilizing the CD-HIT package. The CD-HIT-EST program was employed to eliminate shorter redundant transcripts that were entirely encompassed by other transcripts exhibiting > 90% identity (default setting). The resultant non-redundant clustered transcripts were then classified as unigenes. A total of 184,574 unigenes were predicted from the master/combined assembly **(Table 4**). The total size of unigenes was 19,147,507 bases. The average, maximum, and shortest lengths of unigenes in the libraries were 1037, 19061, and 300 bases, respectively. The size distribution of the raw reads and assembled transcripts from the libraries were characterized, and the data is presented in **Figure 3**.

**Figure 3:**
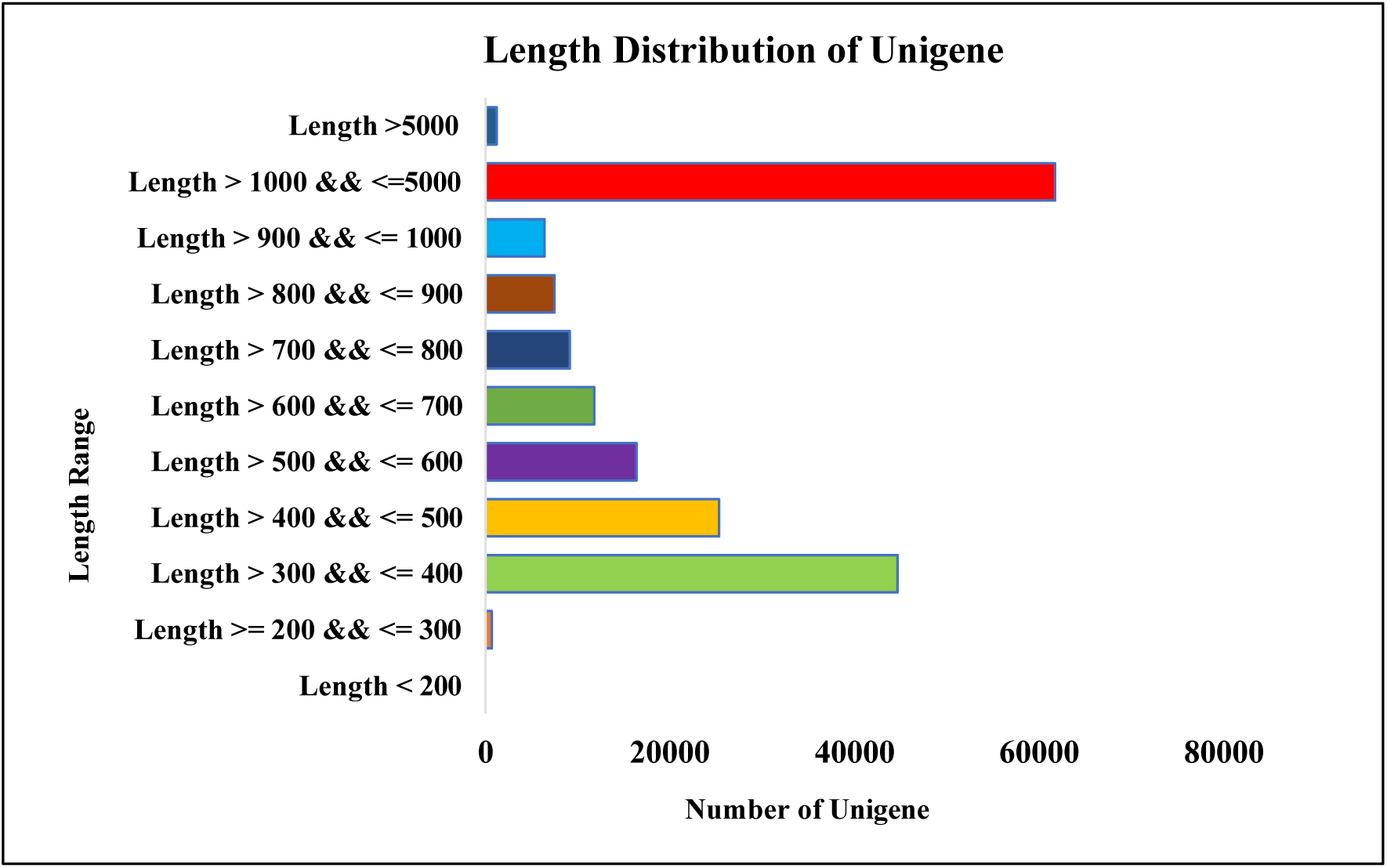
Length distribution of assembled Unigenes. The highest number of Unigenes were observed in the length range of over 1000 to 5000 base pairs, followed by the length range of over 300 to 400 base pairs.

**Table 4:** Unigene assembly statistics predicted from combined assembly.

| Description | Unigenes of *P. chaba* |
| --- | --- |
| Total number of Unigenes | 184574 |
| Total Unigenes size (bp) | 191475070 |
| Average Unigene length (bp) | 1037 |
| Maximum Unigene length (bp) | 19061 |
| Minimum Unigene length (bp) | 300 |
| Unigene N50 (bp) | 1596 |

### CDS obtained from unigenes

CDS were derived from the unigene sequences utilizing Transdecoder with default settings, establishing a minimum encoded protein length of 60 amino acids. This approach identified the segments of unigenes that encode functional proteins, facilitating gene identification and functional annotation. A total of 94,453 CDS nucleotides and a corresponding quantity of CDS proteins were obtained from the identified unigenes. The cumulative size of the CDS nucleotide sequence was 70,981,896 bases, whereas the CDS protein comprised 23,660,632 amino acids (aa). The average, maximum, and minimum lengths of CDS nucleotides were 752, 13,098, and 255 bases, respectively. The subsequent distribution of CDS was conducted based on its length.

### Function annotation of predicted CDS

Functional annotation of the predicted coding sequences (CDS) from the unigenes of three distinct tissues of *P. chaba* was performed through the BLASTX algorithm with an e-value cutoff of 1e-5. The annotation was conducted on the predicted CDS of the *P. chaba* sample by aligning it to the non-redundant protein database of NCBI using BLASTX, with an e-value threshold of less than 1e-5. The BLASTX analysis statistics for the predicted CDS, indicate 80,328 with BLAST hits and 14,125 without, from 94,453 CDS sequences. The species distribution assessment revealed that the predominant hits were associated with *Aristolochia fimbriata*, followed by *Cinnamomum micranthum* (**Figure 4**). All CDS sequences were concurrently analyzed for similarity against Uniprot, KOG, and Pfam utilizing BLASTX with an e-value cutoff of 1e-5. The outcomes of the similarity search across all databases are shown in **Table 5**. KOG classification is an approach for categorizing CDS into functional categories based on orthologous relationships. The KOG analysis demonstrated that the most enriched KOG categories in *P. chaba* were "Signal transduction mechanisms (T)," "General function prediction only (R)," and "post-translational modification, protein turnover, chaperones (O)" (**Figure 5**). Pfam analysis is an approach utilized to detect and categorize protein domains and families within a set of sequences, including transcripts or genes. In the Pfam analysis, the predominant domains discovered in *P. chaba* were "Pkinase Tyr: Protein tyrosine kinase," followed by the "PPR_2 domain" (**Figure 6**).

**Figure 4:**
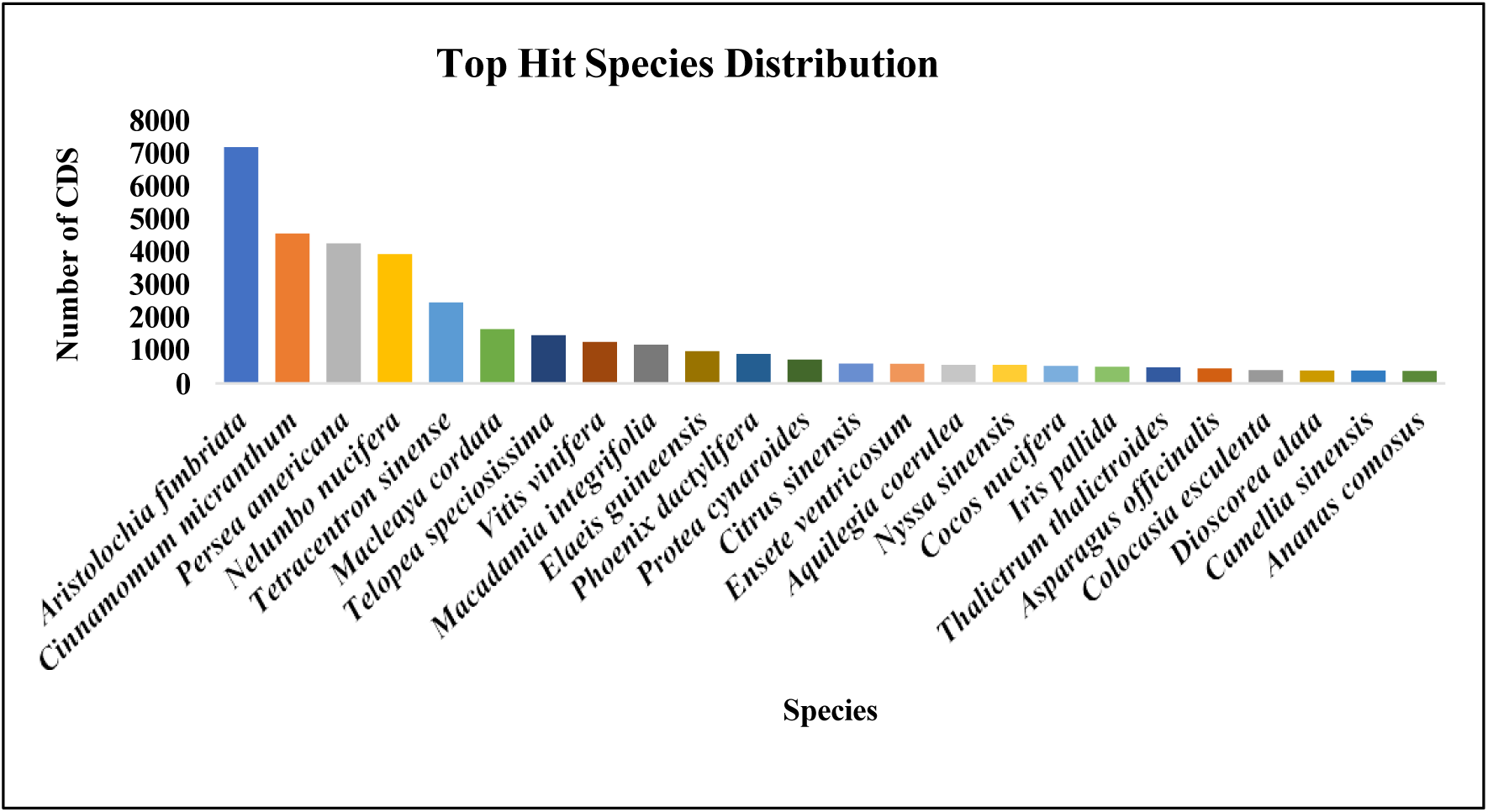
Top species distribution of *Piper chaba* coding sequences (CDS) based on BLAST hits.

**Figure 5:**
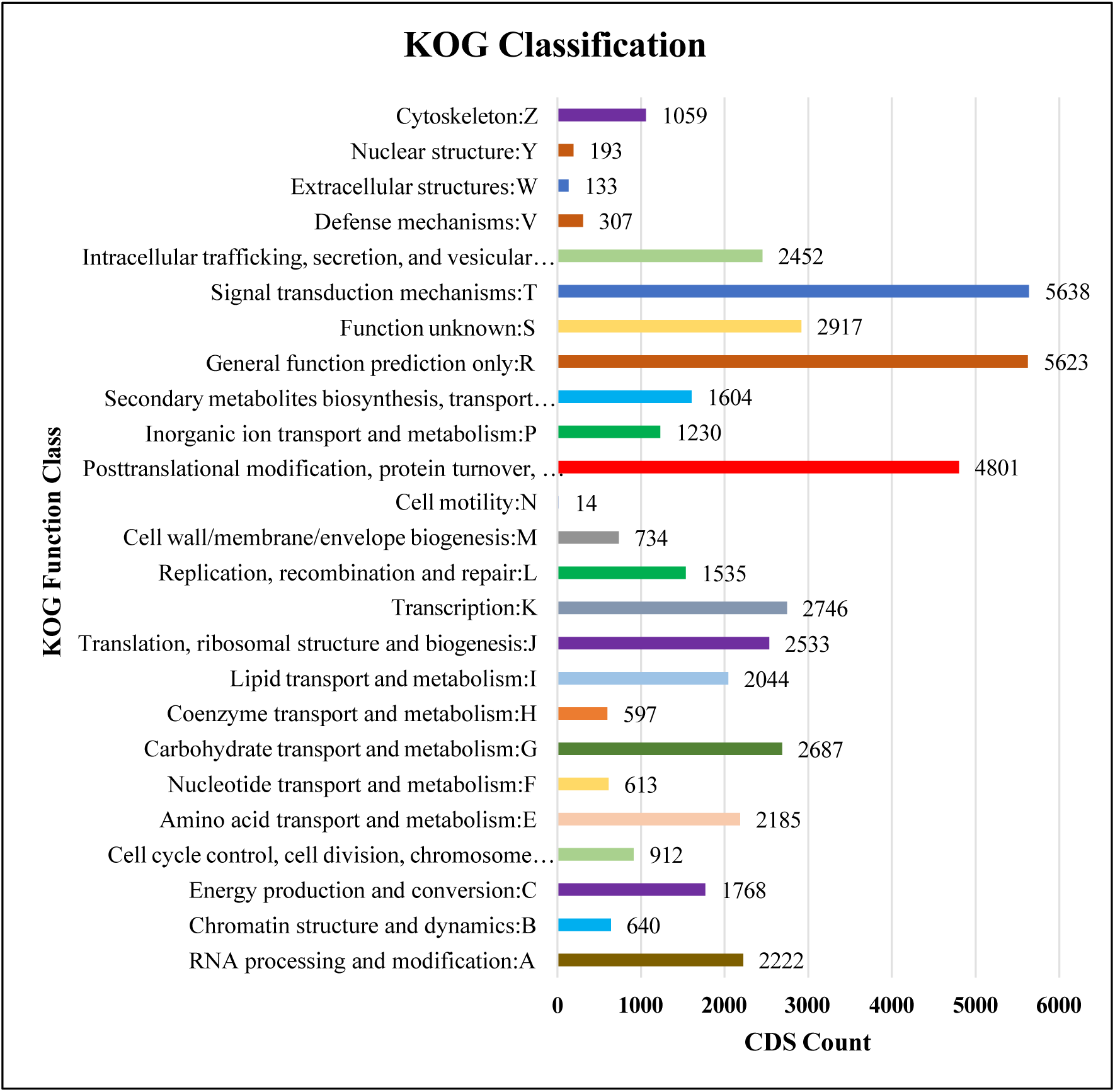
Functional annotation and classification of predicted *Piper chaba* coding sequences (CDS) into 26 KOG functional categories.

**Figure 6:**
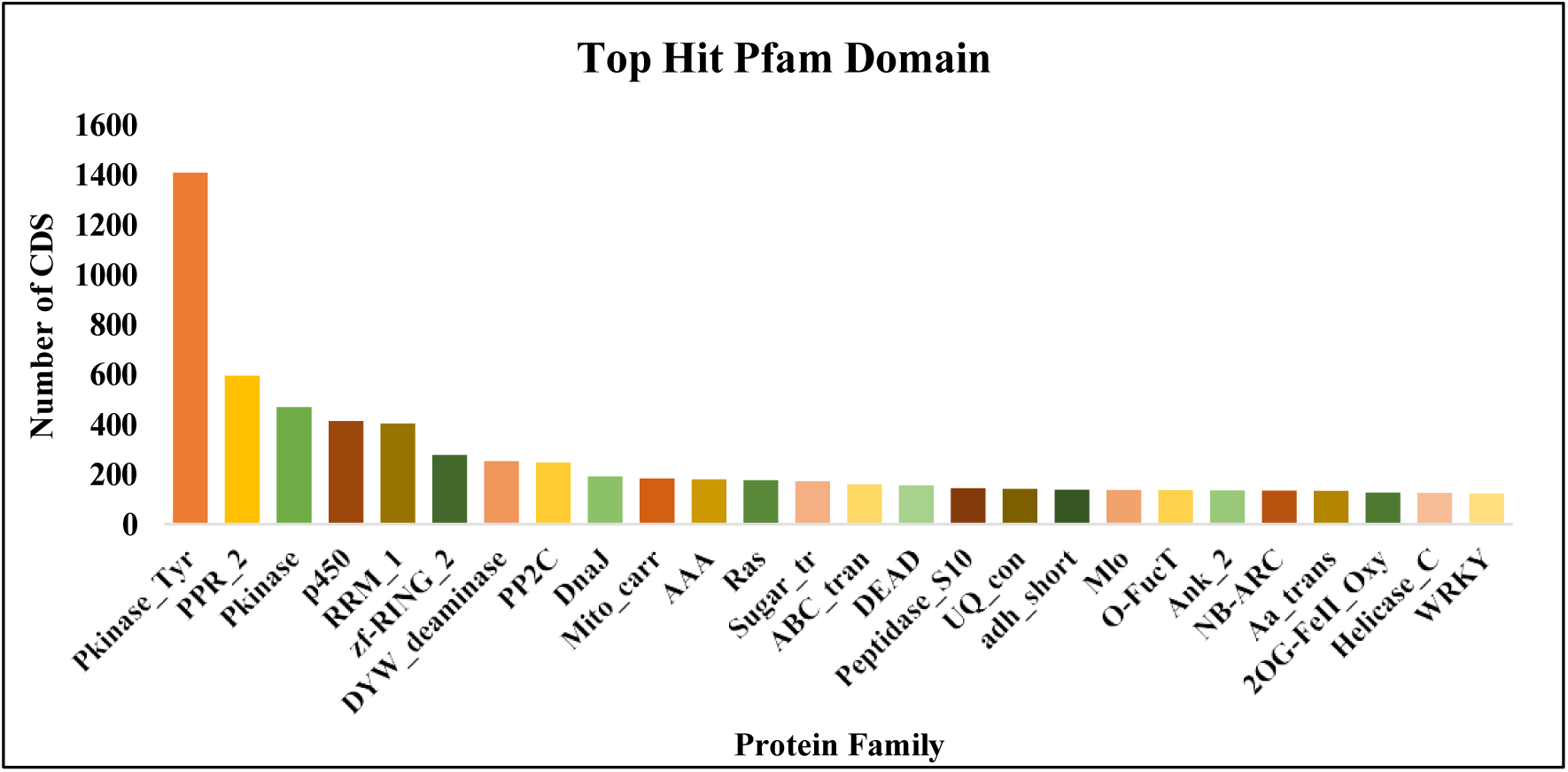
Top Pfam domains identified in *Piper chaba* coding sequences (CDS).

**Table 5:**
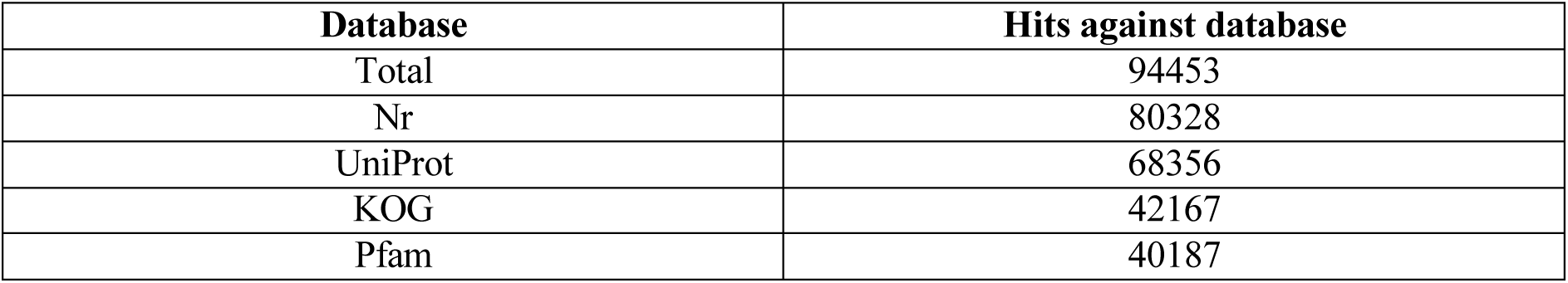
BLAST analysis statistics across protein databases.

| Database | Hits against database |
| --- | --- |
| Total | 94453 |
| Nr | 80328 |
| UniProt | 68356 |
| KOG | 42167 |
| Pfam | 40187 |

A Venn diagram is a visual tool employed to illustrate relationships among several sets of data. It illustrated the similarities and differences among the evaluated databases using overlapping circles. In **Figure 7**, the Venn diagram illustrates that 28,127 CDS were distributed throughout all four databases.

**Figure 7:**
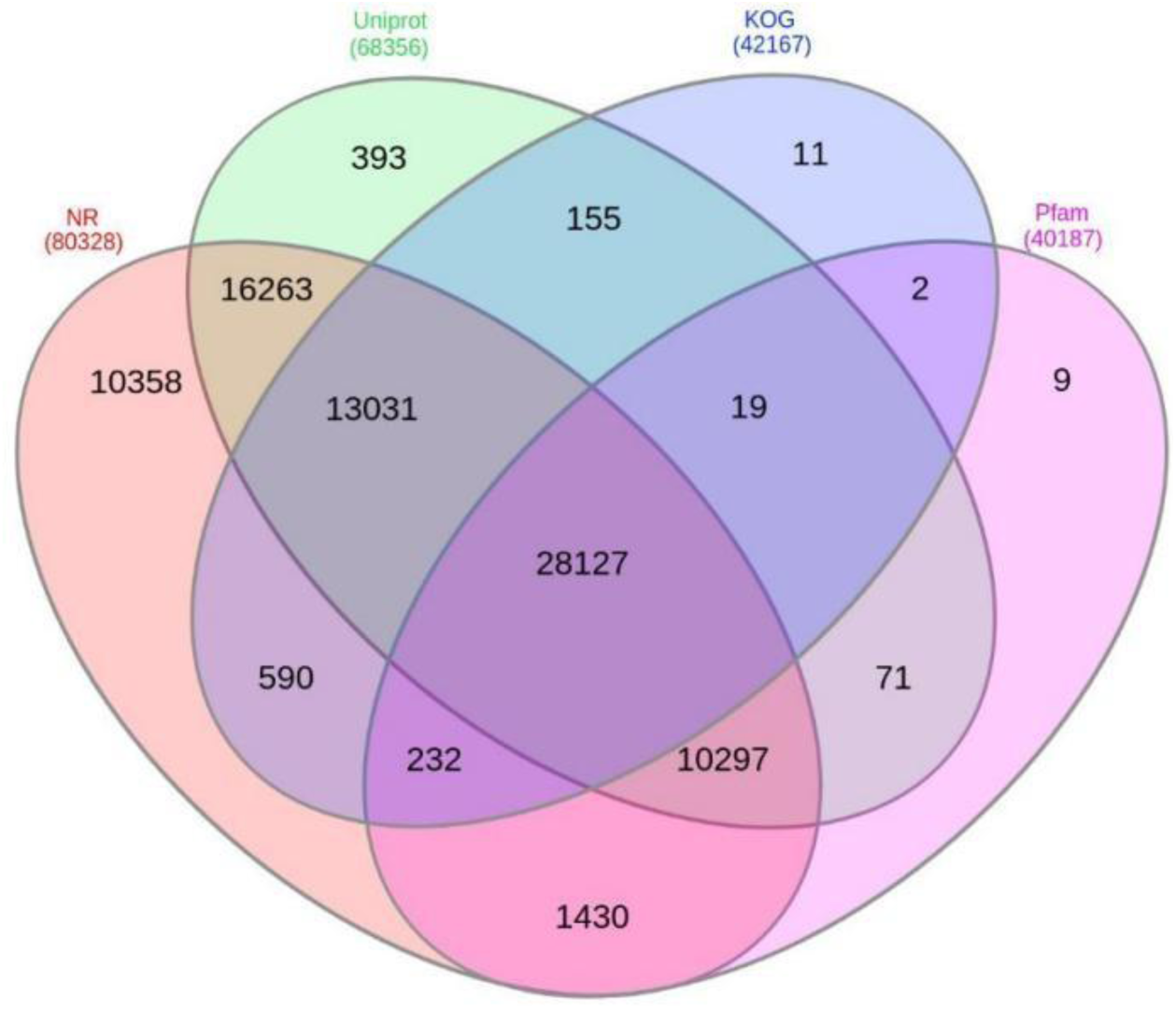
Venn diagram showing the common CDS in NR, Uniport, KOG, and Pfam databases.

### GO sequence distribution of NR annotated CDS

A total of 15,458 CDS were assigned at least one GO term, indicating that a single CDS may possess several GO terms. The Gene Ontology (GO) category is allocated as follows: 10,236 coding sequences (CDS) were designated for Biological Process, 8,195 CDS for Cellular Components, and 12,139 CDS for Molecular Function **(Figure 8**).

**Figure 8:**
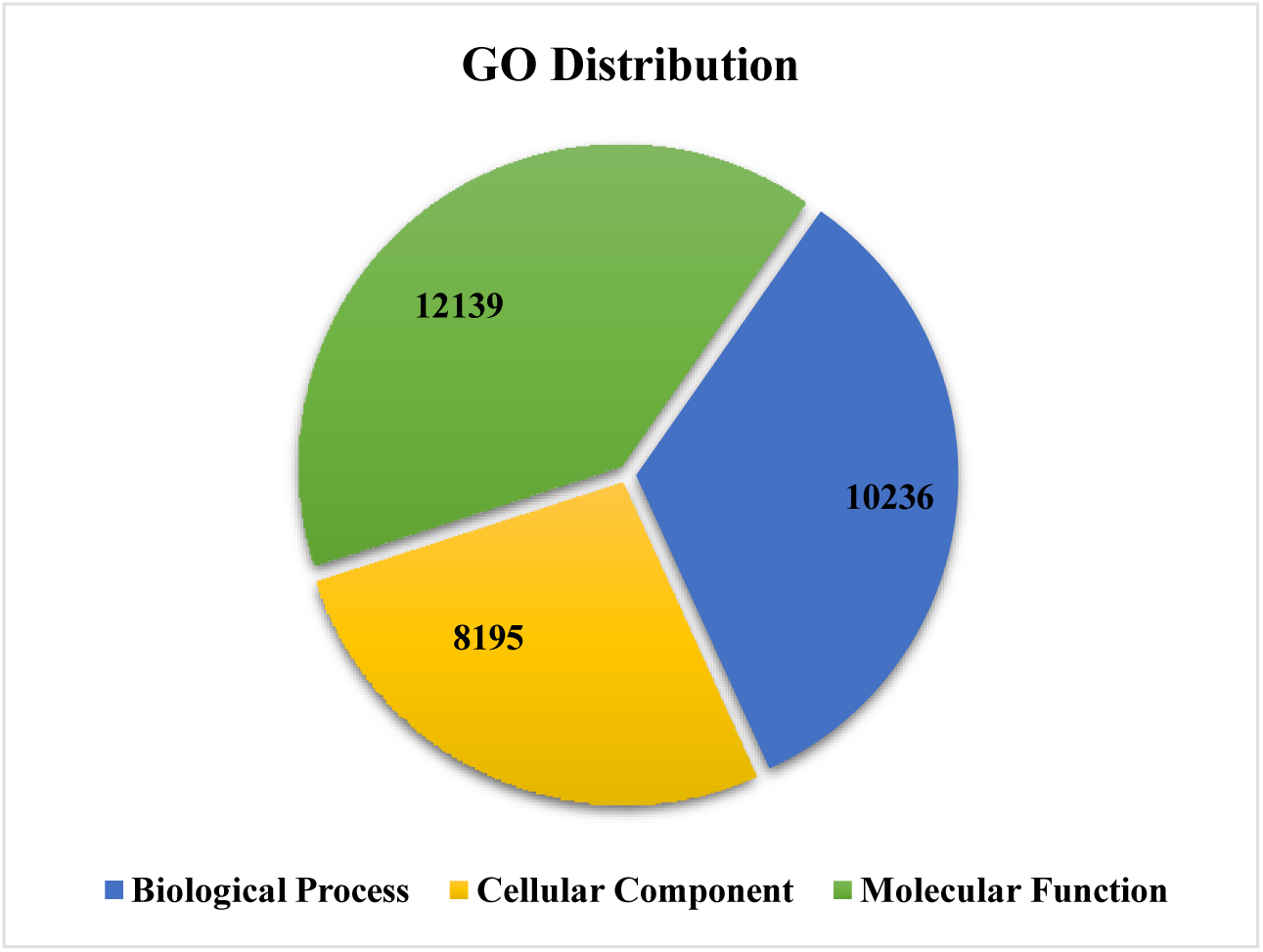
Gene Ontology (GO) classification of *Piper chaba* coding sequences: distribution across Biological Process (BP), Molecular Function (MF), and Cellular Component (CC).

### Transcription Factor (TF) identification

Transcription factors are essential for the synthesis of secondary metabolites. All anticipated CDS were evaluated for similarity against the plant transcription factor database utilizing BLASTX, with an e-value threshold of 1e-5. Among the 58 total hits, the most prevalent transcription factor discovered was bHLH (basic Helix-Loop-Helix), followed by the NAC family. **Figure 9** illustrates the transcripts categorized by main transcription factor families.

**Figure 9:**
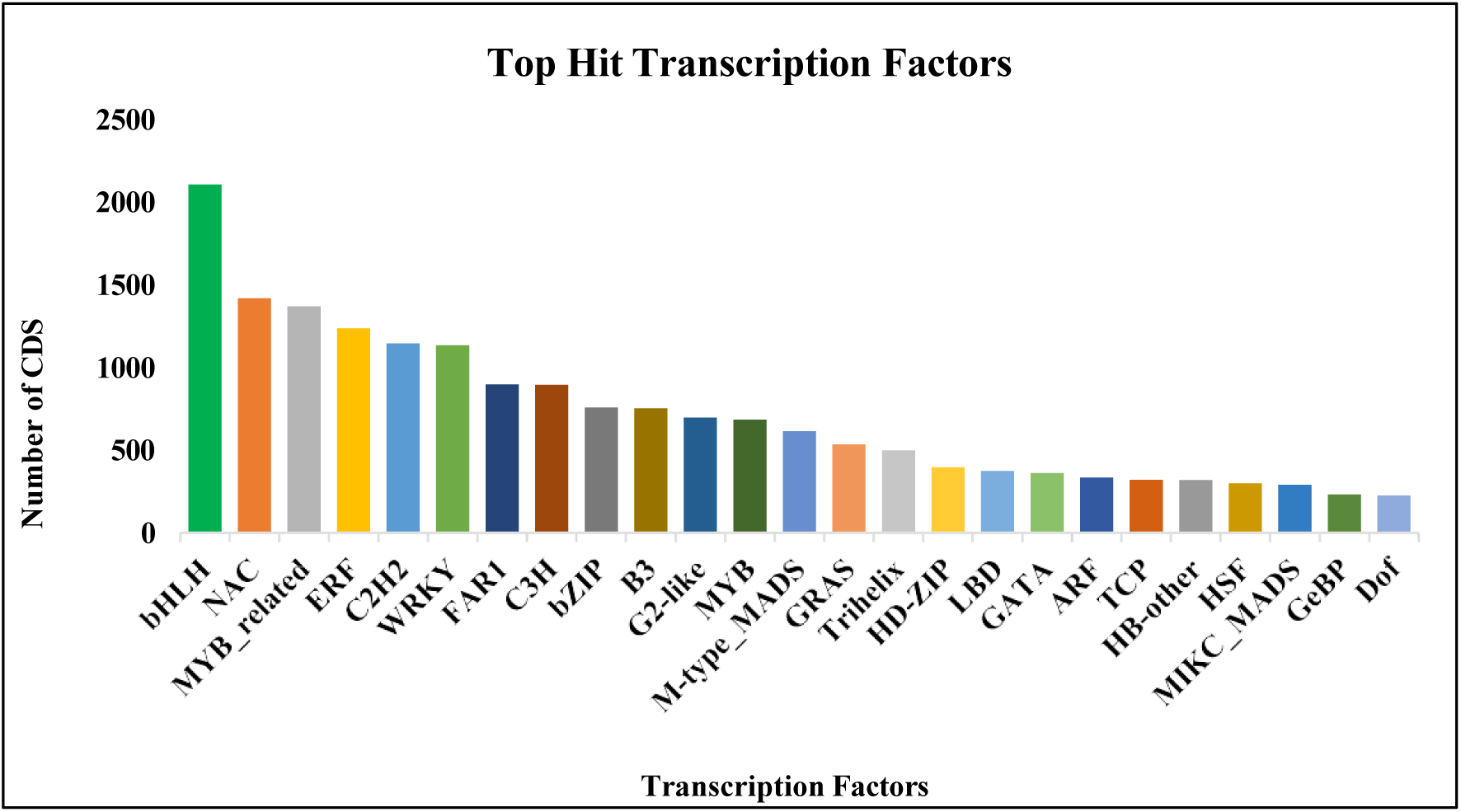
Classification of major transcription factor (TF) families in *Piper chaba* coding sequences (CDS).

### Functional characterization and pathway analysis of CDS using KEGG

Ortholog assignment and the mapping of the coding sequences to biological pathways were conducted utilizing the KEGG Automatic Annotation Server (KAAS). All the CDS were evaluated against the KEGG database utilizing BLASTX with a threshold bit-score value of 60 (default). Of the total 94,453 CDS, 5,876 were aligned with the KEGG database, indicating metabolic pathways of essential macromolecules like carbohydrates, lipids, cofactors, vitamins, amino acids, nucleotides, terpenoids, and polyketides. Genetic information processing includes transcription, translation, folding, sorting, degradation, replication, repair, and information processing in viruses. Environmental information processing includes membrane transport, signal transduction, signaling molecules, and their interactions. The transcripts also indicated the genes associated with metabolism, environmental information processing, genetic information processing, and cellular functions as illustrated in **Figure 10**.

**Figure 10:**
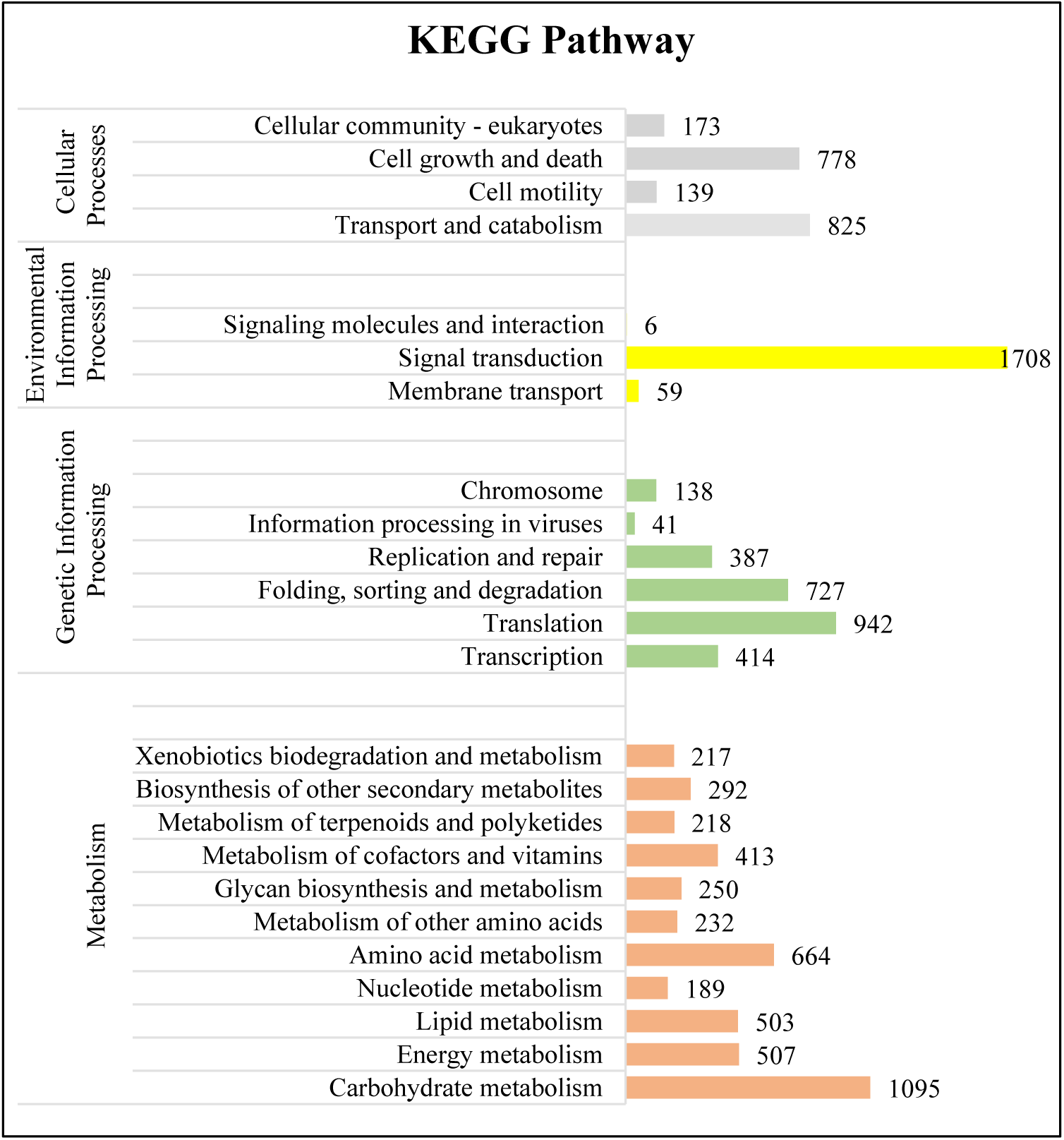
KEGG pathway-based functional annotation and classification of predicted *Piper chaba* coding sequences (CDS).

### Piperine biosynthetic pathway-related genes

In the piperine biosynthetic pathway of *P. chaba*, the thirteen genes were identified including Phenylalanine ammonia-lyase-like protein (Unigene_64981_CDS_71273), Coumarate--CoA ligase (Unigene_51783_CDS_64888), Arogenate dehydratase (Unigene_101207_CDS_576), Cinnamate-4-hydroxylase (Unigene_160753_CDS_38358), Aminotransferase (Unigene_104210_CDS_2524), Cytochrome P450 family (Unigene_113463_CDS_9672), Glycosyl transferase (Unigene_100256_CDS_86), P-Courmaroyl Co – A (Unigene_30201_CDS_50615), 4-Coumarate – Co A ligase (Unigene_51783_CDS_64887), Shikimate hydroxycinnamoyl transferase (Unigene_16628_CDS_39877), Piperic acid Co A ligase (Unigene_140631_CDS_26957), Primary amine oxidase (Unigene_154287_CDS_34411) and Piperamide synthase/BADH acyltransferase (Unigene_147284_CDS_30376). Among the total identified genes, some were associated with lysine metabolism, while others were associated with the phenylpropanoid pathway **(Figure 11).**

**Figure 11:**
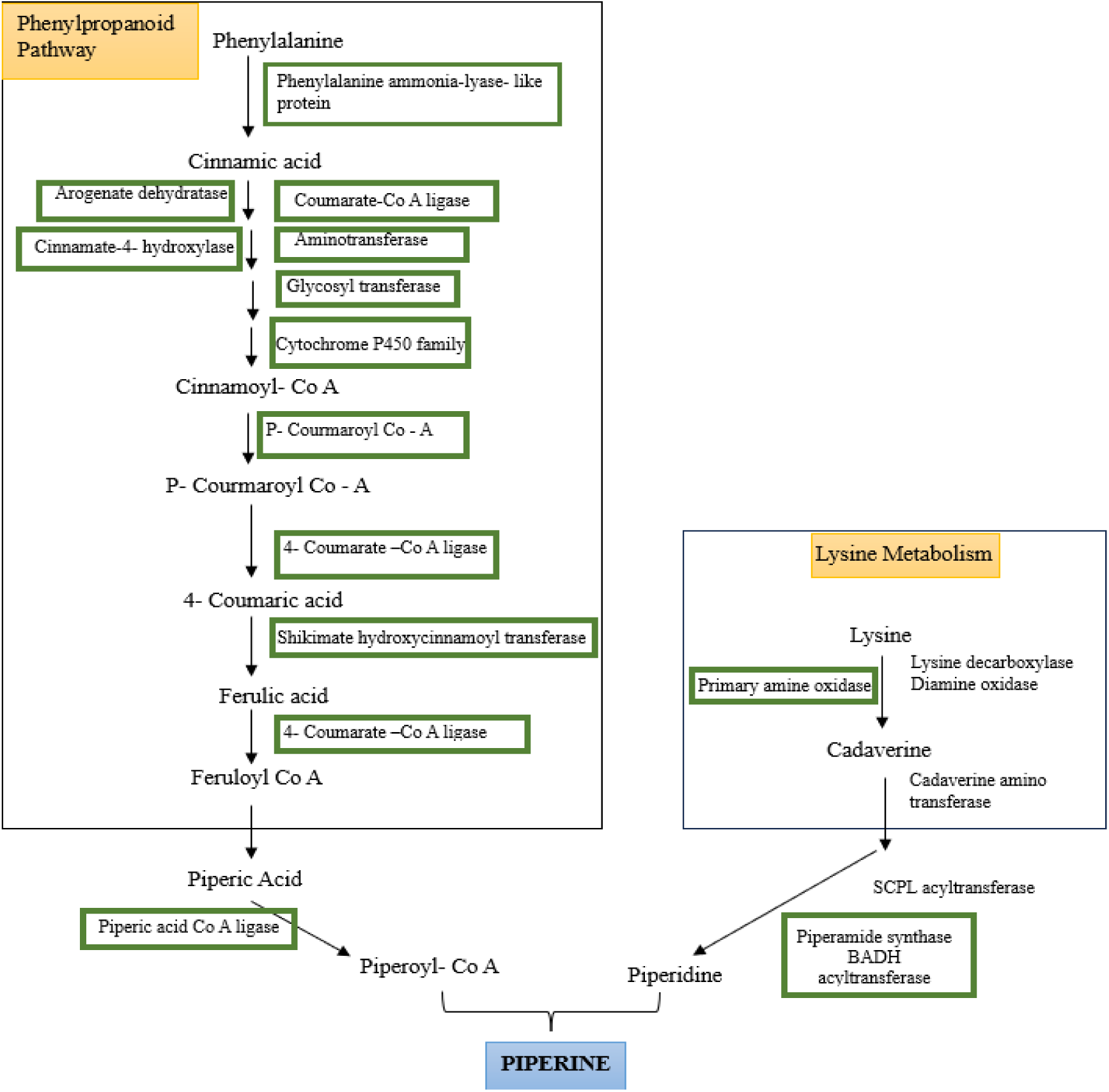
A schematic representation of the key genes involved in the piperine biosynthesis pathway, as identified through the transcriptomic analysis of *Piper chaba*. Genes detected in this study are highlighted within green boxes. The identified genes include **Phenylalanine ammonia-lyase-like protein** (*Unigene_64981_CDS_71273*), **Coumarate-CoA ligase** (*Unigene_51783_CDS_64888*), **Arogenate dehydratase**(*Unigene_101207_CDS_576*), **Cinnamate-4-hydroxylase** (*Unigene_160753_CDS_38358*), **Aminotransferase** (*Unigene_104210_CDS_2524*), **Cytochrome P450 family** (*Unigene_113463_CDS_9672*), **Glycosyl transferase** (*Unigene_100256_CDS_86*), **p-Coumaroyl-CoA** (*Unigene_30201_CDS_50615*), **4-Coumarate-CoA ligase** (*Unigene_51783_CDS_64887*), **Shikimate hydroxycinnamoyl transferase** (*Unigene_16628_CDS_39877*), **Piperic acid-CoA ligase** (*Unigene_140631_CDS_26957*), **Primary amine oxidase** (*Unigene_154287_CDS_34411*), and **Piperamide synthase/BADH acyltransferase** (*Unigene_147284_CDS_30376*).

### Identification of SSR (Simple Sequence Repeats)

Simple sequence repeats (SSRs), also known as microsatellites, are tandem repeats of nucleotide motifs ranging from 2 to 6 base pairs in length. They exhibit significant polymorphism and are universally found in all known genomes. Consequently, SSRs were discovered in the CDS sequences from each sample utilizing the MISA Perl script. Identification was conducted using the following criteria: di-nucleotide patterns must occur a minimum of six times, while tri-, tetra-, penta-, and hexa-nucleotide patterns must appear at least five times. **Table 6** presents the descriptive parameters for the discovered SSR. A total of 94,453 sequences were analyzed, resulting in the identification of 5,050 SSRs. A total of 5,050 SSRs were distributed in unit sizes as follows: 709 dinucleotides, 4,257 trinucleotides, 44 tetranucleotides, 33 pentanucleotides, and 7 hexanucleotides. There are 286 SSRs involved in compound formation. 403 CDS sequences exhibiting multiple SSRs.

**Table 6:** SSR distribution in the master transcriptome assembly.

| Description | Count |
| --- | --- |
| Total number of sequences examined | 94453 |
| Total size of examined sequences (bp) | 70981896 |
| Total number of identified SSRs | 5050 |
| Number of SSR-containing sequences | 4591 |
| Number of sequences containing more than 1 SSR | 403 |
| Number of SSRs present in compound formation | 286 |
| Dinucleotide | 709 |
| Trinucleotide | 4257 |
| Tetranucleotide | 44 |
| Pentanucleotide | 33 |
| Hexanucleotide | 7 |

### Gene expression analysis and validation

To investigate gene expression related to piperine biosynthesis, key genes were selected based on their involvement in molecular functions and biological processes. Ubiquitin C (UbC) was used as an internal control, while eight target genes, including Farnesyl pyrophosphate synthase, Serine-glyoxylate aminotransferase, UDP-glycosyltransferase, Cytochrome P450, Phytoene synthase, Piperic acid synthase, Coumarate-CoA ligase, and Glycosyl transferase, were analyzed. A comprehensive overview of these genes and their functions is provided in **Table 7**.

**Table 7:** Selected genes for validation through qRT-PCR.

| S. No. | CDS | Name of Genes | Function of Genes |
| --- | --- | --- | --- |
| 1. | Unigene_78358_CDS_80420 | Ubiquitin C | Internal control gene |
| 2. | Unigene_88997_CDS_88054 | Farnesyl pyrophosphate synthase | Key enzyme in isoprenoid biosynthetic pathway |
| 3. | Unigene_93637_CDS_90424 | Serine-glyoxylate aminotransferase | Role in the serine assimilation pathway and photorespiration |
| 4. | Unigene_17935_CDS_43258 | UDP-glycosyltransferase | Biosynthesis of natural compounds such as flavonoids, terpenoids |
| 5. | Unigene_134546_CDS_22123 | Cytochrome P450 | Regulate the biosynthesis and catabolism of plant hormones like auxins, cytokinins, and gibberellins. |
| 6. | Unigene_73189_CDS_77315 | Phytoene synthase | Carotenoid biosynthesis pathway |
| 7. | Unigene_81681_CDS_82640 | Piperic acid synthase | Biosynthetic pathway of piperine |
| 8. | Unigene_51783_CDS_64887 | Coumarate-CoA ligase | Role in the biosynthesis of piperine by activating intermediate acids to corresponding CoA |
| 9. | Unigene_100256_CDS_86 | Glycosyl transferase | Catalyzed the biosynthesis of complex carbohydrates |

For primer design, the Integrated DNA Technologies (IDT) online tool was utilized to generate primers with optimized amplicon length, melting temperature, and GC content. **Table 8** details the forward and reverse primer sequences used for gene validation.

**Table 8:**
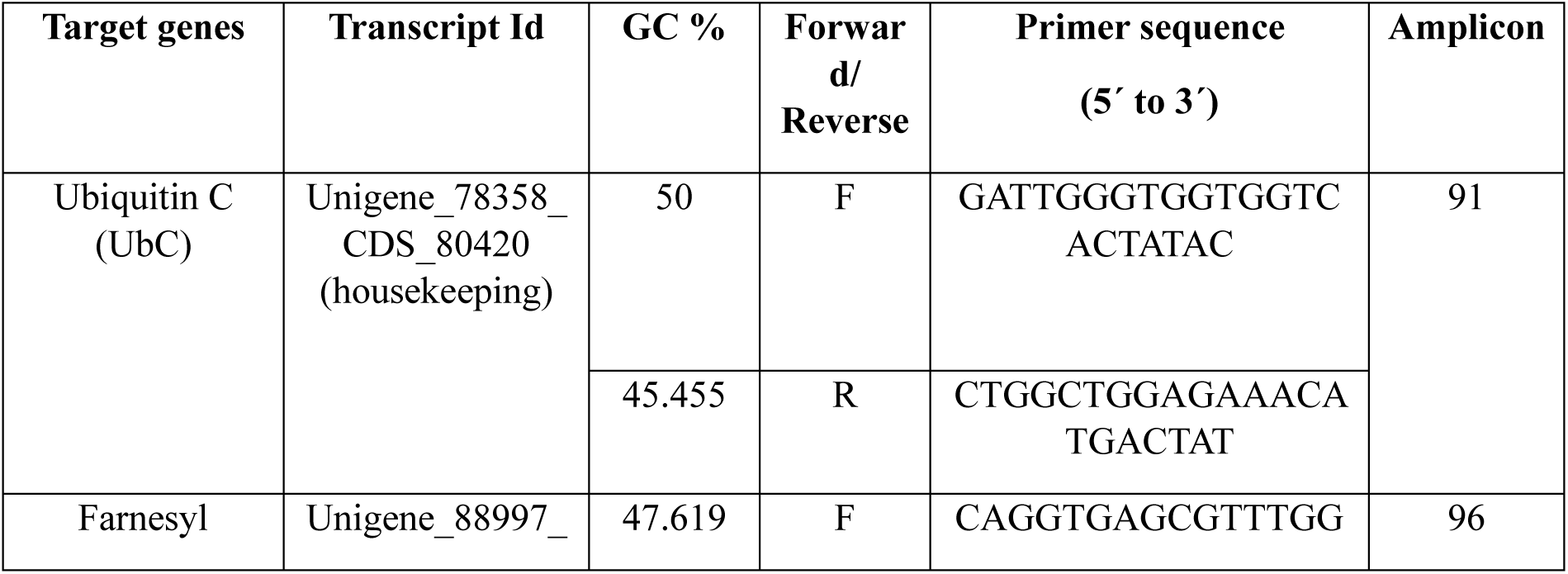
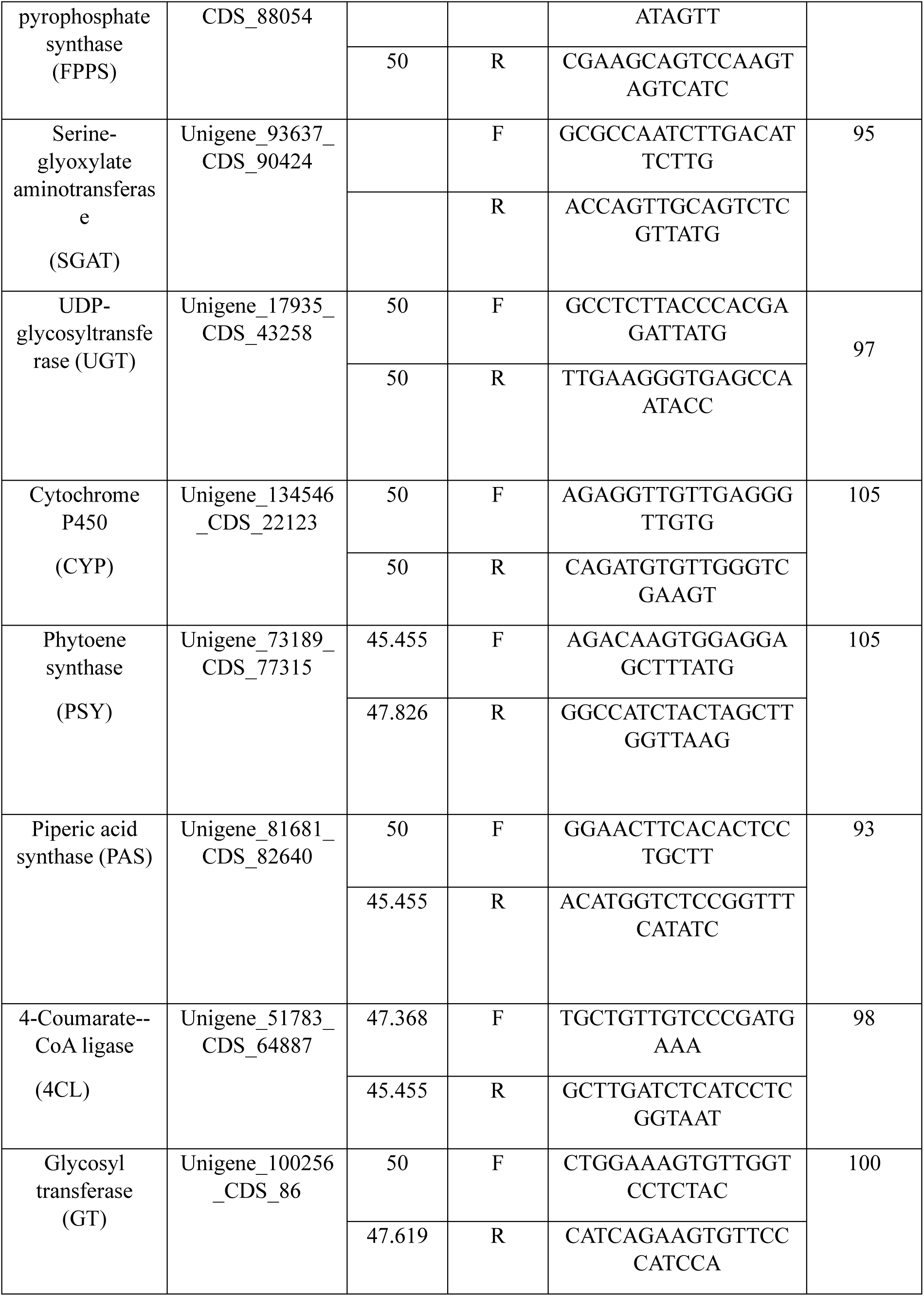
qRT-PCR primers for validation of selected genes in *Piper chaba*.

RNA was isolated using the PureLink RNA Mini Kit, followed by qualitative and quantitative analysis. First-strand cDNA synthesis was carried out using the R2D 1st-Strand cDNA Synthesis Kit. The quality of synthesized cDNA was confirmed using gene-specific primers. The PCR products were separated using 1.2% agarose gel electrophoresis.

Quantitative gene expression analysis was conducted through qRT-PCR to validate differentially expressed genes identified in leaf-vs-root and spike-vs-root comparisons. qRT-PCR reactions included a control lacking the template, and UbC served as the internal reference gene (**Table 9**). Relative transcript levels were assessed using the 2^−ΔΔCt^ method. Gene expression profiles (log2 fold change) were statistically analyzed (± SE) and presented in **Figures 12 and 13**, ensuring accuracy through three biological and technical replicates.

**Figure 12:**
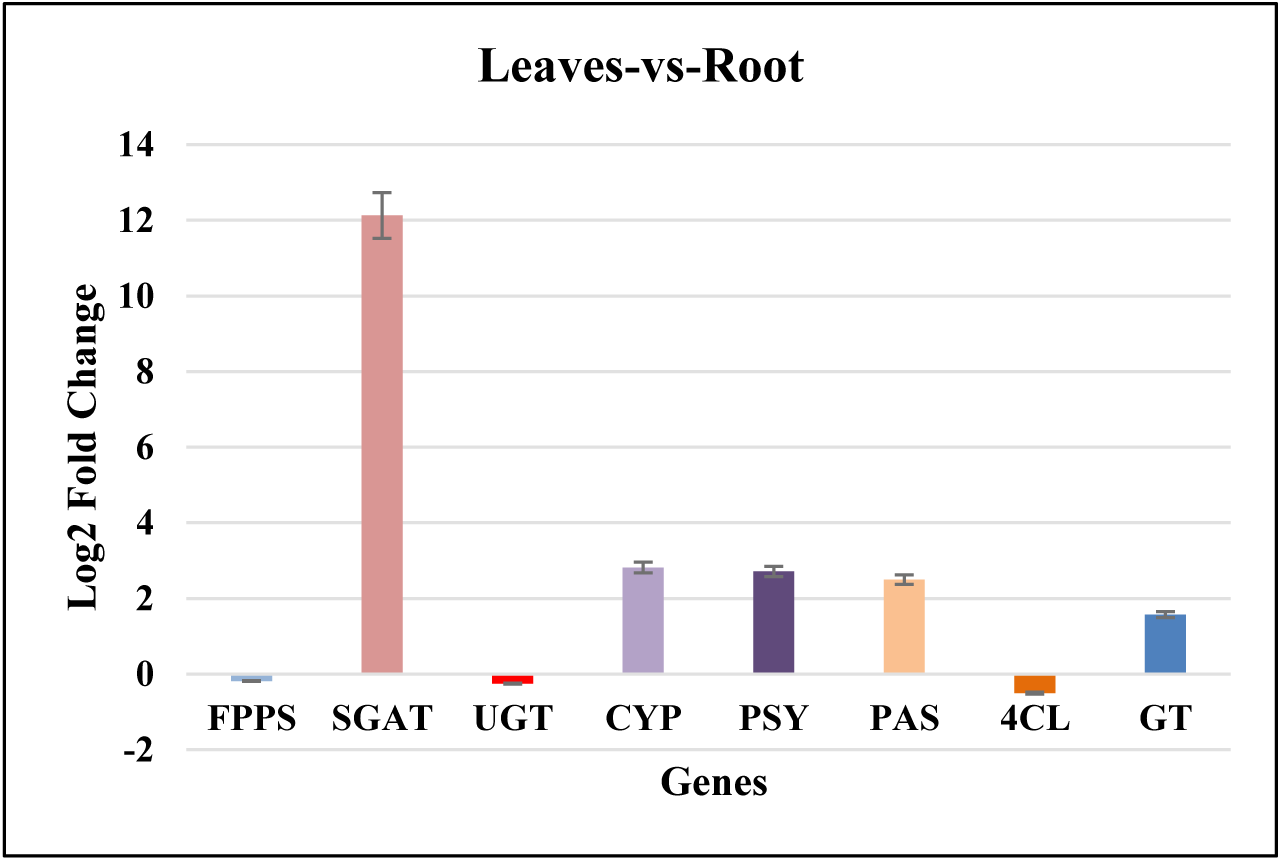
Graphical representation of fold change in selected genes for Leaves vs. Root comparison.

**Figure 13:**
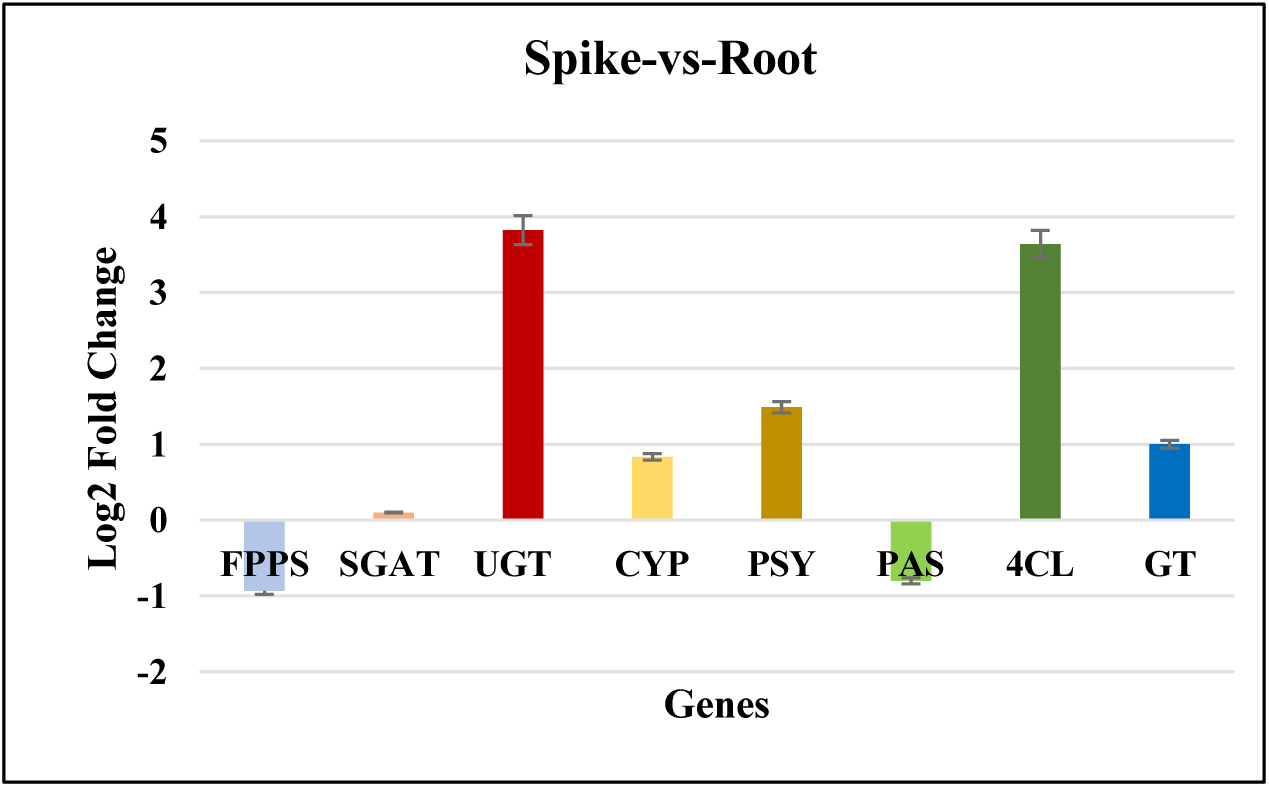
Graphical representation of fold change in selected genes for Spike vs. Root comparison.

**Table 9:** Log2 fold change in differential expression of selected genes.

| S. No. | Gene | log2 fold change (Root-vs- Leaves) | log2 fold change (Root-vs- spike) |
| --- | --- | --- | --- |
| 1. | Ubiquitin C (UbC) | 0 | 0 |
| 2. | Farnesyl pyrophosphate synthase (FPPS) | -0.18686 | -0.93207 |
| 3. | Serine-glyoxylate aminotransferase (SGAT) | 12.12573 | 0.098595 |
| 4. | UDP-glycosyltransferase (UGT) | -0.25232 | 3.824167 |
| 5. | Cytochrome P450 (CYP) | 2.815387 | 0.833674 |
| 6. | Phytoene synthase (PSY) | 2.713209 | 1.487527 |
| 7. | Piperic acid synthase (PAS) | 2.496661 | -0.80129 |
| 8. | 4-Coumarate--CoA ligase (4CL) | -0.50464 | 3.638918 |
| 9. | Glycosyl transferase (GT) | 1.569168 | 0.999972 |

## Discussion

Transcriptome analyses of the root, flower, and fruit of *Piper nigrum, P. longum,* and *P. retrofractum* had been conducted independently, providing preliminary genetic insights into the biosynthesis of piperine (Hu et al. 2015; Moreira et al. 2017; Khew et al. 2020; George et al. 2022; Yadav et al. 2023; Meechuen et al. 2023; Prasad et al. 2025). The limited genetic resources for *P. chaba* have made molecular investigations of regulatory mechanisms difficult. In the present research, the transcriptomes of *P. chaba* from root, leaves, and spike were *de novo* sequenced and assembled. Gene expression profiling related to the piperine biosynthesis pathway was conducted, and comparisons of organ-specific transcriptomes and metabolite accumulations were made. The findings elucidated the molecular mechanisms behind the synthesis of piperine in *P. chaba*. Overall, this research provided important genetic resources for *P. chaba* research, facilitating future molecular research and potential improvements in herbal quality by targeting the metabolic pathways related to the production of natural compounds.

The biosynthesis pathway of piperine remained partially understood, with several genes implicated in the process still to be fully characterized. The majority of research on piperine biosynthesis has been performed in *Piper nigrum* (Schnabel et al. 2021b; Yadav et al. 2023; Lv et al. 2024). Recent studies have outlined a suggested biosynthetic pathway whereby piperoyl-CoA and piperidine are derived from phenylalanine and lysine, respectively (Jin et al. 2020; Cotinguiba et al. 2022; Lv et al. 2023). The BAHD-acyltransferase enzyme, pipBAHD1, catalyzes the reaction between piperoyl-CoA and piperidine to produce piperine (Schnabel et al. 2021a). Recent investigations indicate that piperoyl-CoA is synthesized from piperic acid by the action of piperoyl-CoA ligase (Jin et al. 2020; Dantu et al. 2021; Jackel et al. 2022; IP et al. 2024).

A literature review indicated that insufficient focus has been placed on the spatiotemporal distribution of piperine and its precursors in *P. chaba* but several studies conducted on other species of piper provided hypothetical pathways of piperine biosynthesis (Hu et al, 2019; Schnabel et al. 2021a; Lv et al. 2023). The next-generation sequencing investigation of the pathways involved in the production of natural compounds like piperine in *P. chaba* would facilitate the identification of more natural products for the advancement of new pharmaceuticals and the manipulation of plant processes. The transcriptome offers a cost-effective method for sequence determination, enhancing the efficiency and speed of gene discovery. *P. chaba* is a significant therapeutic plant; yet, transcriptomic, and genomic data were not present at the NCBI database.

This study conducted a transcriptome analysis of leaves, root, and spike of *P. chaba*, yielding high-quality transcriptomic data through the Illumina NovaSeq 6000 platform. Total RNA was isolated from the collected plant samples with the Alexgen Total RNA Extraction Kit. A 500-1000 ng aliquot of isolated RNA was utilized to construct a paired-end sequencing library using the KAPA mRNA HyperPrep Kit for Illumina. The concentration, yield, and absorbance of extended libraries were evaluated using the TapeStation (Agilent 4150). The library profiles of *P. chaba* leaves, roots, and spikes exhibited a singular or average peak, signifying a high-quality library. The sequencing generated ∼6.74, ∼11.18, and ∼10.55 GB of high-quality reads from the leaves, roots, and spikes of *P. chaba*, respectively. A total of 2,11,99,256 leaves, 3,51,63,291 spike, and 3,31,70,294 root raw reads were obtained through the Illumina platform. *De novo* master assembled from high-quality reads of *P. chaba* samples was constructed by using Trinity software at default parameters (kmer 25). 228,481 transcripts were analyzed in the *de novo* assembly of *P. chaba*. Transcripts undergo additional processing with CD-HIT software to predict unigenes. Using Transdecoder with the default parameters, CDS were inferred from the unigene sequences. A total of 184,574 unigenes and 94,453 CDS were predicted. The raw reads were subsequently uploaded to the Sequence Read Archive (SRA) of the NCBI database under accession numbers SRR28673946, SRR28673947, and SRR28673948 for the leaves, root, and spike of *P. chaba*, respectively. To visualize the transcriptome comprehensively, analyzed CDS were functionally annotated and classified. The functional annotation was performed on the predicted coding sequences of the plant sample by aligning the CDS with the non-redundant protein database of NCBI utilizing BLASTX, with a minimum e-value threshold of less than 1e-5. *P. chaba* CDS was aligned with many available protein databases. All CDS were queried against the KOG, Pfam, and UniProt databases using BLASTX. The maximum number of CDS hits with NR databases (80322) followed by Uniport (68356), KOG (42167), and Pfam (40187).

The ten most prevalent Pfam domains were Pkinase Tyr, PPR 2, Pkinase, p450, RRM 1, zf-RING 2, DYW deaminase, PP2C, DnaJ, and Mito carr. The distribution of top-hit species indicated a similarity between *P. chaba* and *Aristolochia fimbriata, Cinnamomum micranthum, Persea americana, Nelumbo nucifera, Tetracentron sinense, Macleaya cordata, Telopea speciosissima, Vitis vinifera*, and *Macadamia integrifolia*. The species distribution in the annotation showed the evolutionary relationship of *P. chaba* with other species.

Transcription factors modulate gene expression in response to external and internal signals, activating or repressing downstream genes. During the annotation, the evaluation of the transcriptome data of *P. chaba* revealed several transcription factors from diverse families. These families are essential for regulating various secondary metabolites and playing a crucial role in piperine production. C3H transcription factors are part of the zinc finger motif family, which facilitates interactions with other molecules (Liu et al. 2020). Certain variables influenced pollen generation, secondary metabolism regulation, flower growth, protein binding, and shoot apical meristem development. Overall, 21,350 CDS were annotated with the transcription factors (TFs) database and categorized into 58 transcription factor families. In our study, the predominant transcription factor families involved in synthesis were identified, with the bHLH family showing the maximum number of CDS hits at 2107. This was succeeded by the MYB-related family with 2,732 CDS hits, NAC with 2,695 CDS hits, ERF with 2,320 CDS hits, C2H2 with 1,957 CDS hits, C3H with 1,792 CDS hits, WRKY with 1,758 CDS hits, B3 with 1,566 CDS hits, FAR1 with 1,564 CDS hits, and bZIP with 1,429 CDS hits, all of which demonstrated high expression levels. The basic helix-loop-helix (bHLH) family constituted the most extensive group in this dataset. These transcription factors are essential in modulating plant growth and development, encompassing light signal transduction, the formation of specialized cells, and hormone responses. They also participate in secondary metabolism, particularly under stress conditions and in response to abiotic and biotic resistance. The MYB family constitutes one of the largest and most varied transcription factor families in plants (Cao et al. 2020). They are essential in regulating cell differentiation, developmental processes, and responses to environmental stresses. Their function in the production of secondary metabolites, hormone signaling, and growth regulation was essential. This dataset indicated that bHLH, MYB-related, NAC, and ERF transcription factors predominate, underscoring their essential function in regulating growth, development, and stress responses in the organism. The elevated number of bHLH and MYB-related factors indicated that these pathways were essential for metabolic regulation and environmental adaptability. The WRKY and ERF families illustrated the critical roles of pathogen defense and stress tolerance, while FAR1 and bZIP transcription factors emphasized the importance of light response and developmental control.

SSRs, also known as microsatellites, are tandem repeats of nucleotide motifs ranging from 2 to 6 base pairs in length. They exhibit significant polymorphism and are found in all known genomes. A total of 5050 SSR markers were discovered from the transcriptome data of *P. chaba*. In total, the identified SSRs included 709 di-nucleotide, 4257 tri-nucleotide, 44 tetra-nucleotide, 33 pentanucleotide, and 7 hexanucleotide repeats, with tri-nucleotides being the most prevalent and hexanucleotides the least common in *P. chaba*. A total of 2034 SSR markers were discovered from 19899 ESTs in *Catharanthus roseus* by the development of EST-based SSR markers. The efficacy of these markers was evidenced by their high repeatability, the number of bands that may be scored for each marker, their transferability, and their polymorphism (Mishra et al. 2011).

The predicted roles of CDS had been determined by GO assignments. The GO mapping categorized the predicted CDS into three primary domains: Biological Process with 10,236 CDS, Cellular Component with 8,195 CDS, and Molecular Function with 12,139 CDS. The CDS were compared with the KEGG database to further understand the functions of several metabolic pathways in *P. chaba*. Orthologs were assigned, and the coding sequences were mapped to biological pathways utilizing the KAAS. A preset threshold bit-score of 60 was employed in BLASTX to compare all coding sequences against the KEGG database. The distribution of 5,876 CDS was classified into four main categories and several subcategories. The mapped CDS also represented genes associated with environmental information processing, metabolic functions, genetic information processing, and cellular processes. The enzyme with designated functions, comprising 4580 CDS inside a metabolic pathway in KEGG, was illustrated. Among these CDS, 1095 were related to glucose metabolism, whereas 664 were assigned to amino acid metabolism. 507 and 503 identified CDS were associated with energy and lipid metabolism respectively. The highest number of CDS associated with signal transduction was 1708. In total 217 coding sequences were associated with the biosynthesis of terpenoids and polyketides. Additionally, 21 CDS were implicated in the metabolism of tropane, piperidine, and pyridine alkaloids.

Transcriptome study of the leaves, root, and spike discovered gene families related to the biosynthesis pathway of piperine and other secondary metabolites. Thirteen genes involved in piperine biosynthesis were identified, including Phenylalanine ammonia-lyase-like protein, Coumarate-CoA ligase, Arogenate dehydratase, Cinnamate-4-hydroxylase, Aminotransferase, Cytochrome P450 family, Glycosyl transferase, P-Coumaroyl Co-A, 4-Coumarate-CoA ligase, Shikimate hydroxycinnamoyl transferase, Piperic acid CoA ligase, Primary amine oxidase, and Piperamide synthase/BADH acyltransferase.

To validate the Illumina sequencing expression profiles, nine genes including Ubiquitin C (UbC), Farnesyl pyrophosphate synthase (FPPS), Serine--glyoxylate aminotransferase (SGAT), UDP-glycosyltransferase (UGT), Cytochrome P450 (CYP), Phytoene synthase (PSY), Piperic acid synthase (PAS), 4-Coumarate--CoA ligase (4CL), Glycosyl transferase (GT) were randomly selected for qRT-PCR analysis to confirm further their expression levels, derived from differentially expressed genes discovered by comparative transcriptome study of the leaves, roots, and spikes of *P. chaba.* The qRT-PCR results showed that the RNA-seq data obtained from this research was reliable. The results of qRT-PCR revealed that serine-glyoxylate aminotransferase is mostly expressed in the leaves. At the same time, UDP-glycosyltransferase and 4-coumarate-CoA ligase demonstrate elevated expression in the spikes, but both were downregulated in the leaves.

This data is an essential reference for forthcoming functional genomics research on *P. chaba* and similar species. Comprehensive phytochemical and molecular investigations would be crucial to identify the genes essential for piperine biosynthesis.

## Conclusion

Unraveling the genetic basis of secondary metabolite biosynthesis is essential for harnessing the medicinal potential of plants. This study provides the first comprehensive transcriptomic analysis of *P. chaba*, offering critical insights into its molecular framework. UPLC-based quantification of piperine across different plant parts confirmed its highest concentration in the spike, emphasizing its pharmacological relevance. High-quality RNA sequencing and *de novo* transcriptome assembly identified 184,574 unigenes and 94,453 coding sequences, which were functionally annotated to elucidate key biosynthetic pathways. Differential gene expression analysis highlighted the transcriptional regulation of piperine biosynthesis, further validated through qRT-PCR. The identification of key transcription factors, such as bHLH and NAC families, along with 5,050 SSR markers, provides valuable genomic resources for future research. By establishing a molecular foundation for *P. chaba*, this study not only enhances our understanding of its genetic and metabolic pathways but also paves the way for its conservation, genetic improvement, and pharmaceutical applications. These findings contribute to the sustainable utilization of *P. chaba*, positioning it as a promising candidate for biotechnological advancements in medicine and agriculture.

## Acknowledgments

The authors sincerely thank the Director, of Dayalbagh Educational Institute, Dayalbagh, Agra, for providing the necessary infrastructure and support for this research. Deeksha Singh acknowledges the financial assistance from **CSIR-UGC**, New Delhi, through the Junior and Senior Research Fellowships.

